# Adaptive value of circadian rhythms in High Arctic Svalbard ptarmigan

**DOI:** 10.1101/2020.08.17.254011

**Authors:** Daniel Appenroth, Gabriela C. Wagner, David G. Hazlerigg, Alexander C. West

**Affiliations:** Arctic Chronobiology and Physiology, University of Tromsø, Framstredet 42, 9019 Tromsø, Norway; Divisjon for skog og utmark, NIBIO, Holtveien 66, 9016 Tromsø, Norway

**Keywords:** Photoperiodism, Circadian, Seasonal reproduction, Pars tuberalis, Svalbard ptarmigan, the Arctic

## Abstract

The arctic archipelago of Svalbard (74 to 81° North) experiences extended periods of uninterrupted daylight in summer and uninterrupted darkness in winter. Species native to Svalbard display no daily rhythms in behaviour or physiology during these seasons, leading to the view that circadian rhythms may be redundant in arctic environments [1, 2]. Nevertheless, seasonal changes in the physiology and behaviour of arctic species rely on photoperiodic synchronisation to the solar year. Since this phenomenon is generally circadian-based in temperate species, we investigated if this might be a preserved aspect of arctic temporal organisation.

Here, we demonstrate the involvement of the circadian clock in the seasonal photoperiodic response of the Svalbard ptarmigan (*Lagopus muta hyperborea*), the world’s northernmost resident bird species. First, we show the persistence of rhythmic clock gene expression under constant conditions within the mediobasal hypothalamus and pars tuberalis, the key tissues in the seasonal neuroendocrine cascade. We then employ a “sliding skeleton photoperiod” protocol, revealing that the driving force behind seasonal biology of the Svalbard ptarmigan is rhythmic sensitivity to light, a feature that depends on a functioning circadian rhythm. Our results suggest that the unusual selective pressure of the Arctic relaxes the adaptive value of the circadian clock for organisation of daily activity patterns, whilst preserving its importance for seasonal synchronisation. Thus, our data simultaneously reconnects circadian rhythms to life in the Arctic and establishes a universal principle of evolutionary value for circadian rhythms in seasonal biology.

## RESULTS AND DISCUSSION

### The rhythmic expression of circadian clock genes in the mediobasal hypothalamus and pars tuberalis of Svalbard ptarmigan persists under constant light

Svalbard ptarmigan (Figure 1A) show diurnal behaviour patterns under daily light-dark cycles, but are behaviourally arrhythmic in constant light conditions (Figure 1B) [1, 3]. These data, and similar findings in Svalbard reindeer [2, 4], suggest that circadian rhythms are redundant in the high arctic habitat of Svalbard. The Svalbard ptarmigan, however, uses photoperiod to time seasonal changes in its physiology [3, 5–7], and a vast collection of data supports the role of the circadian rhythm in photoperiodic timekeeping [8–17]. Thus, although the ptarmigan circadian rhythm may not be required for maintaining daily synchrony, it may play an essential role in photoperiodic responses.

**Figure 1.**
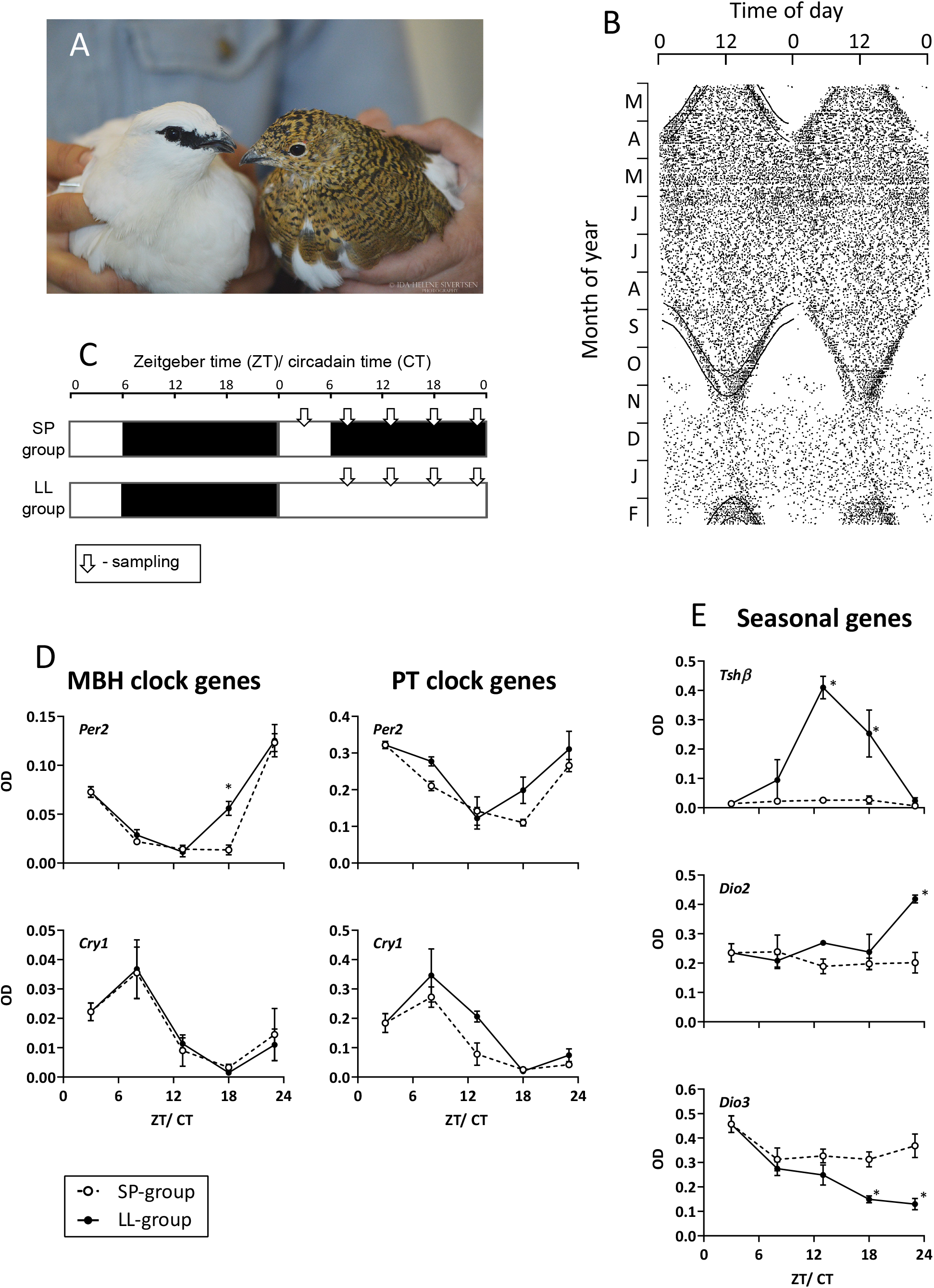
Persistence of circadian rhythmicity in the pars tuberalis and mediobasal hypothalamus of Svalbard ptarmigan. (A) Svalbard ptarmigan in different plumages. On the right a male in white winter plumage and on the left a female in brown summer plumage (© Ida-Helene Sivertsen). (B) Double plotted actogram of a Svalbard ptarmigan in Svalbard over one year (March to February). Thick lines indicate on- and offset of civil twilight and thin lines indicate sunrise and sunset. Redrawn using data from Reierth *et al*. (1998) [1]. (C) Experimental design. Birds entrained to SP (6L:18D) either remained in SP or were transferred directly into LL. Samplings are indicated by arrows and are given in Zeitgeber time (ZT) or circadian time for the LL-group (CT). Both groups were sampled at ZT/ CT 8, 13, 18 and 23. The SP-group was additionally sampled at ZT 3 (ZT 3 was used as initial point for plotting both group, but was omitted from statistical analysis). (D) Gene expression for *Per2* and *Cry1* in the MBH and PT between the SP- (dashed line) and LL-group (solid line). Data is displayed as mean optical densities (OD) ± SEM. Asterisks indicate significant differences between the groups at a given ZT/ CT (p < 0.05 by by Sidak’s *post hoc* test). (E) Gene expression for *Tshβ, Dio2* and *Dio3* in the PT and MBH between the SP- (dashed line) and LL-group (solid line). Gene expression was measured by radioactive *in situ* hybridization and is displayed as mean OD ± SEM. Asterisks indicate significant differences between the groups at a given ZT/ CT (p < 0.05 by by Sidak’s *post hoc* test).

We first used radioactive *in situ* hybridization to examine the transcriptional regulation of circadian genes *Cry1* and *Per2* within the mediobasal hypothalamus (MBH) and pars tuberalis (PT) of the pituitary gland, since these sites control the seasonal neuroendocrine response in other gallinaceous species [18–20]. Our results showed that both genes were strongly rhythmic under short photoperiod (SP, L6:D18) and displayed negligible changes in their expression patterns following transfer to constant light (LL) (Figure 1C and 1D). Hence, core elements of the avian circadian clock show persistent endogenous rhythmicity in key photoperiodic response tissues.

In temperate and tropical bird species [18–23] the seasonal reproductive response depends on photoperiodic control of thyrotropin beta subunit (*Tshβ*) expression in the PT and consequent thyrotropin receptor-mediated changes in MBH function, exemplified by changes in the expression of the thyroid hormone deiodinase genes, *Dio2* and *Dio3*. In the Svalbard ptarmigan *Tshβ* expression in the PT was continuously suppressed under SP, and transfer to LL strongly induced *Tshβ* expression, which peaked 13 h after lights-on (CT13) (p < 0.0001 compared to SP control birds by Sidak’s *post hoc* test) (Figure 1E) before falling back to SP levels by 23 h after lights-on (CT23). Within the MBH, transfer to LL significantly induced the expression of *Dio2* by CT23 (p = 0.0011 by Sidak’s *post hoc* test), and suppressed the expression of *Dio3* by CT18 (p = 0.0085 by Sidak’s *post hoc* test) (Figure 1E). These data show that the temporal dynamics of the “first long day” photoperiodic neuroendocrine response is highly conserved between Svalbard ptarmigan and their relatives from temperate latitudes, i.e. Japanese quail (*Coturnix japonica*) [18].

### A sliding skeleton photoperiod triggers the long-day seasonal response in Svalbard ptarmigan

In 1936, Erwin Bünning proposed that photoperiodic sensitivity depends on a circadian rhythm in light sensitivity [24]. A wealth of data supports this hypothesis, confirming that short light-pulses given during a so-called ‘photoinducible phase’ are sufficient to drive long-day seasonal response [8–14]. In other words, it is not the cumulative duration of light exposure that triggers a long-day response, but the coincidence of light with an endogenously defined circadian phase.

To test the involvement of circadian rhythms in photoperiodic sensitivity of Svalbard ptarmigan, we exposed our birds to either extended SP, an increasing continuous photoperiod (IP) or a sliding skeleton photoperiod (SkP). The SkP-group mimics the extending range of the IP-group, but maintains the same cumulative hours of light in a 24-h period as in the SP-group (Figure 2A). Expression of a long-day phenotype in the SkP would therefore demonstrate a circadian rhythm in photoperiodic sensitivity in Svalbard ptarmigan. To track the development of a long-day phenotype we monitored activity, body mass, food intake and plasma testosterone; variables that are all under photoperiodic control in Svalbard ptarmigan [5–7, 25].

**Figure 2.**
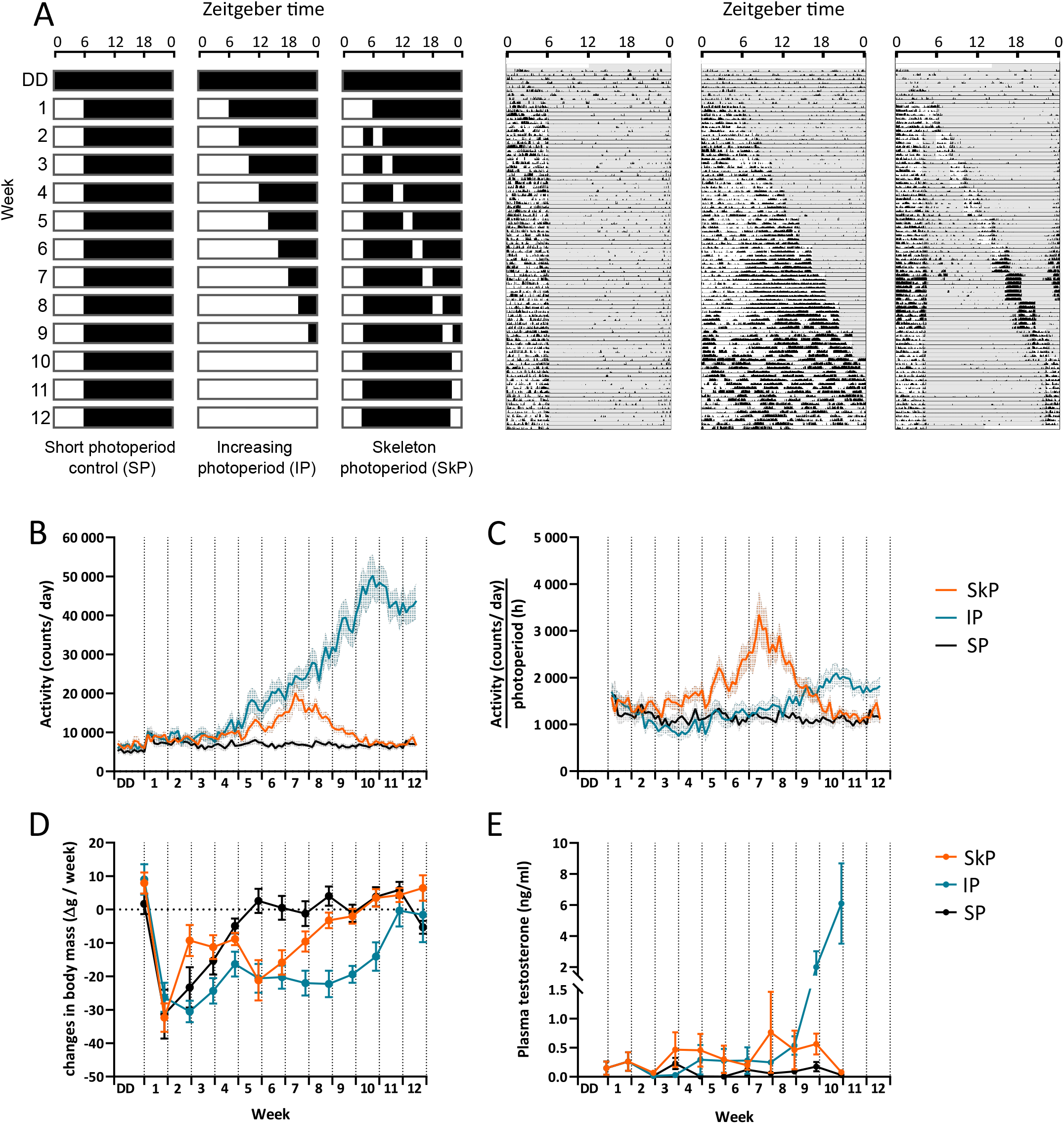
Physiological and endocrine responses to increasing photoperiod and a sliding skeleton photoperiod. (A) Experimental design. All bird were initially transferred from DD to SP (6L:18D), which marked the start of the experiment. Birds of the SP-group remained under SP for 12 weeks. Birds of the IP-group were subjected to a stepwise increase in photoperiod by extending the lights-off signal by two hours every week. The light-period of the SkP-group was split into two blocks of light at week 2. The long 4h light-period remained static while the 2-h light-period shifted forwards weekly by two hours. By week 10 the light-period merged again at which point the birds were back to SP but shifted forward by two hours. Representative single-plotted actograms are displayed next to photoperiodic treatments. Grey shading in the actograms indicate periods of darkness. (B) Activity profiles for each group measured as count/day and displayed as means ± SEM. (C) Activity profiles presented as counts/ day divided by the hours of light. Data is displayed as means ± SEM. (C) Changes in body mass measured as grams gained or lost from one week to another. Data is presented as means ± SEM (D) Plasma testosterone of male birds measured as ng/ml and displayed as means ± SEM.

We observed a strong diurnal activity preference within all the groups (Figure 2A). The activity profile of the SP-group went unchanged throughout the entire experiment; however, both the IP and SkP groups increased their activity between weeks 5 and 7 (Figure 2B) (p < 0.05 for all IP vs SP and SkP vs SP between week 5 and 7 by Tukey’s *post hoc* test). Whereas the activity increase in the IP-group within this period was proportional to the increased hours of light, the activity of the SkP-group showed a marked 3-fold increase in intensity within the restricted light-hours (p < 0.05 for all SkP vs SP and SkP vs IP in week 7 by Tukey’s *post hoc* test) (Figure 2C), indicating a photoperiodic stimulation of activity.

Associated with these increases in activity, we observed sustained declines in body mass in both the SkP and IP groups continuing until weeks 9 and 11 respectively (Figure 2D). Food intake was similar between all three groups until week 10 (Figure S1B), suggesting that these responses were a consequence of increased activity resulting in a negative energy balance. Longitudinal assessment of plasma testosterone in male birds showed a clear stimulation in week 10 in the IP-group (p < 0.0001 for IP vs SP and IP vs SkP by Tukey’s *post hoc* test) (Figure 2E), but no statistically significant changes in the other two groups. Hence the intensification of activity in SkP birds and in IP birds prior to week 10 is not a secondary consequence of gonadal changes, but probably reflects photoperiodic induction of pre-breeding territorial behaviour [26, 27]

While the activity level of the IP-group continued to rise throughout the experiment, with maximal activity once the birds experienced LL, the activity of the SkP-group reduced after week 7, returning to SP levels by week 10 (p > 0.05 for SkP vs SP at all points from week 10 onwards by Tukey’s *post hoc* test, Figure 2B and 2C). The reversal of the response in SkP-but not IP-birds probably reflects either a movement of the secondary 2-h block of light beyond the photosensitive phase or a “phase jump” in the entrainment of the circadian rhythm, so that the extended dark interval following the 4-h light-block re-aligns from subjective day to subjective night [28]. Whichever scenario applies, the birds behaved as if experiencing a declining photoperiod after week 7 even though cumulative daily light exposure remained constant. Overall, these results suggest that the Svalbard ptarmigan use a circadian-based system to mediate spring photoperiodic induction of pre-breeding behaviour, with a photoinducible phase some 14 to 16 h after dawn (ZT 14-16).

### A sliding skeleton photoperiod triggers the long-day photoperiodic neuroendocrine cascade in Svalbard ptarmigan

SkP shows a strong photo-stimulatory effect at week 6 where the second (2-h) light-period falls 14 h after the start of the first light-period. We performed a second experiment to determine if these behavioural and physiological responses correspond to classical photoperiodic regulation of the molecular neuroendocrine cascade within the PT/ MBH region. We compared Svalbard ptarmigan under SP to a SkP in which from week 6 onwards the 2-h block of light was held at 14 to 16 h after the start of the 4-h block of light, i.e. to coincide with the photoinducible phase inferred from the previous experiment (Figure 3A). Longitudinal measurements of activity, body mass, food intake and testosterone, were consistent with our previous experiment, and showed a persistent impact of the ZT 14-16 light-period on the development of a summer phenotype (Figure S2). The SkP-group shows increased expression of *Dio2* (p = 0.0024 by unpaired t-test) and decreased expression of *Dio3* (p = 0.0011 by unpaired t-test) (Figure 3B). This indicates that through light-stimulation of the photoinducible phase we were able to elicit the classically described changes in MBH thyroid hormone metabolism in our experimental birds. This cements the role of the circadian clock in photoperiodic sensitivity in the Svalbard ptarmigan. Radioactive *in situ* hybridization analysis for *Tshβ* showed no change between treatments (p = 0.2589 by unpaired t-test). The low *Tshβ* expression in the SkP-group is most likely due to the sampling of the birds at ZT 0.5-1.5 which is comparable with the early light-period in our “first long day” experiment, when *Tshβ* expression is low (Figure 1E). This data additionally suggest comparable circadian phase between the SP and SkP groups.

**Figure 3.**
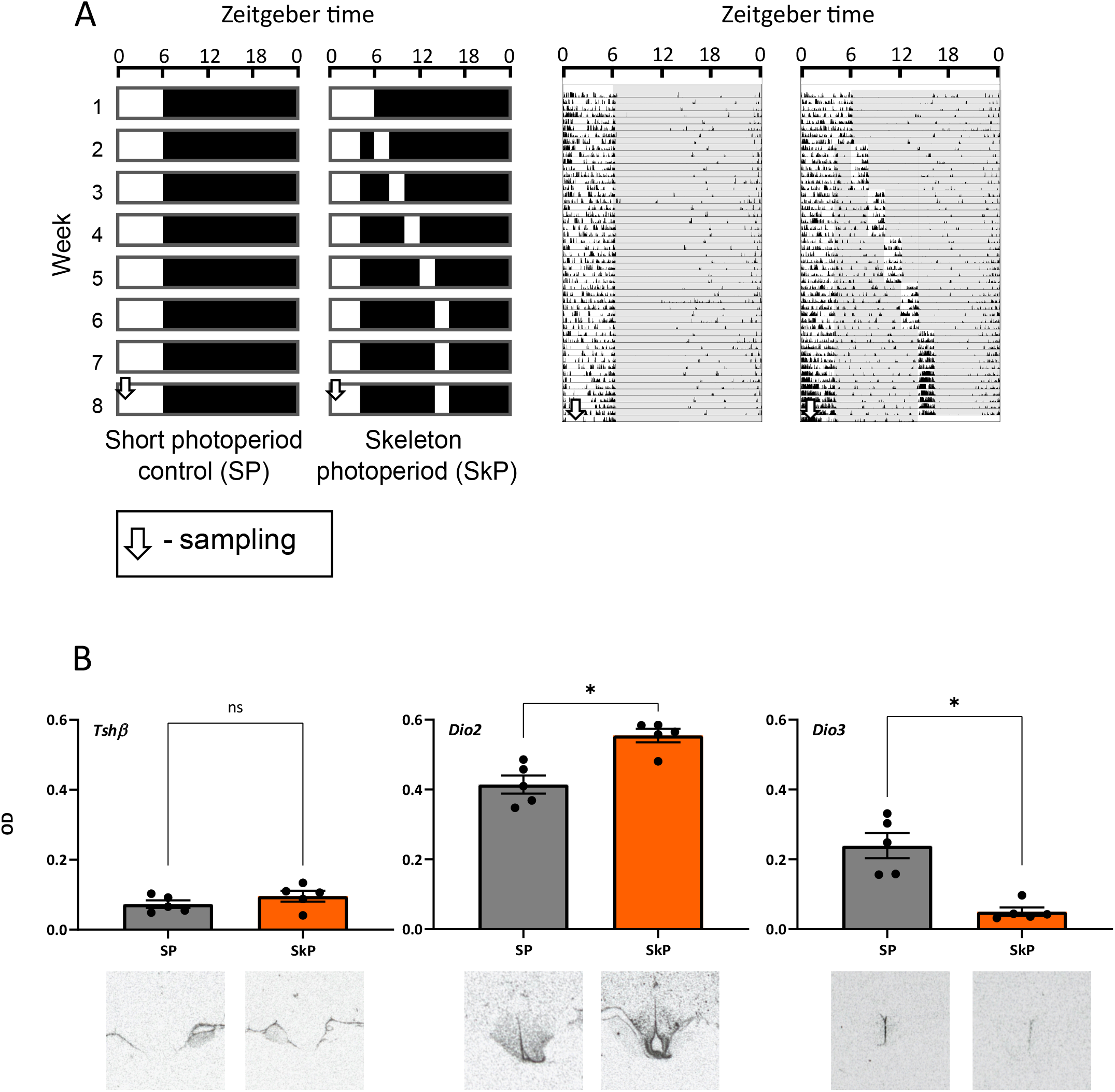
Response of photo-induced genes in the PT and MBH to a skeleton photoperiod. (A) Experimental design. Birds entrained to SP (6L:18D) either remained in SP for 8 weeks or experienced a shifting skeleton photoperiod. The light-period of the SkP-group was split into a 4-h and a 2-h light-period in week 2. The 2-h light-period was weekly shifted backward by two hours until week 6 at which point the light-period remained at ZT 14-16 for three weeks. All birds were sampled in week 8 at ZT 0.5-1.5. Representative single-plotted actograms are displayed next to photoperiodic treatment. Grey shading in the actograms indicate periods of darkness. (B) Gene expression of *Tshβ, Dio2* and *Dio3* in the PT and MBH, measured by *in situ* hybridization. Data is presented as mean optical densities (OD) ± SEM and asterisks indicate significant differences between the groups (p < 0.05 by unpaired t-test). Representative radiographs are displayed under the respective gene and group.

### Adaptive value of circadian rhythms in the Arctic

Overall, our results affirm the universal adaptive role of the circadian rhythm for photoperiodic time measurement. We show that an organism in which daily behavioural rhythms disappear under the midnight sun and during the polar night, maintains a rhythmic molecular clock in tissues essential for seasonal timing and employs a circadian rhythm to set a photoinducible circadian phase.

The unusual phenotypes of the Svalbard ptarmigan and Svalbard reindeer, which shows a similar circadian phenotype [2, 4, 29, 30], reflect the unusual selective pressures of their habitat. Summer and winter on Svalbard provide two wildly contrasting environments for which seasonal synchronisation is essential, particularly of reproductive physiology. Conversely, the daily amplitude of irradiance and temperature are greatly reduced on Svalbard compared to lower latitudes, thus the adaptive value of daily behavioural organisation on Svalbard is likely to be weak, and may even exert counter-adaptive constraints on exploitation of the polar summer (Figure 4). Nevertheless, several reports of sustained circadian behaviour and physiology in other arctic dwellers have emerged, e.g. Arctic ground squirrels [31, 32], bumblebees [33], polar bears [34], copepods [35] and several migrating birds [36–38]. Collectively, these studies suggest that species-specific differences in circadian outputs should be seen as flexible solutions to complex life-history constraints [39]. For the Svalbard ptarmigan, the adaptive solution is to maintain a circadian rhythm to measure photoperiod, but decouple behavioural drive as a circadian output.

**Figure 4.**
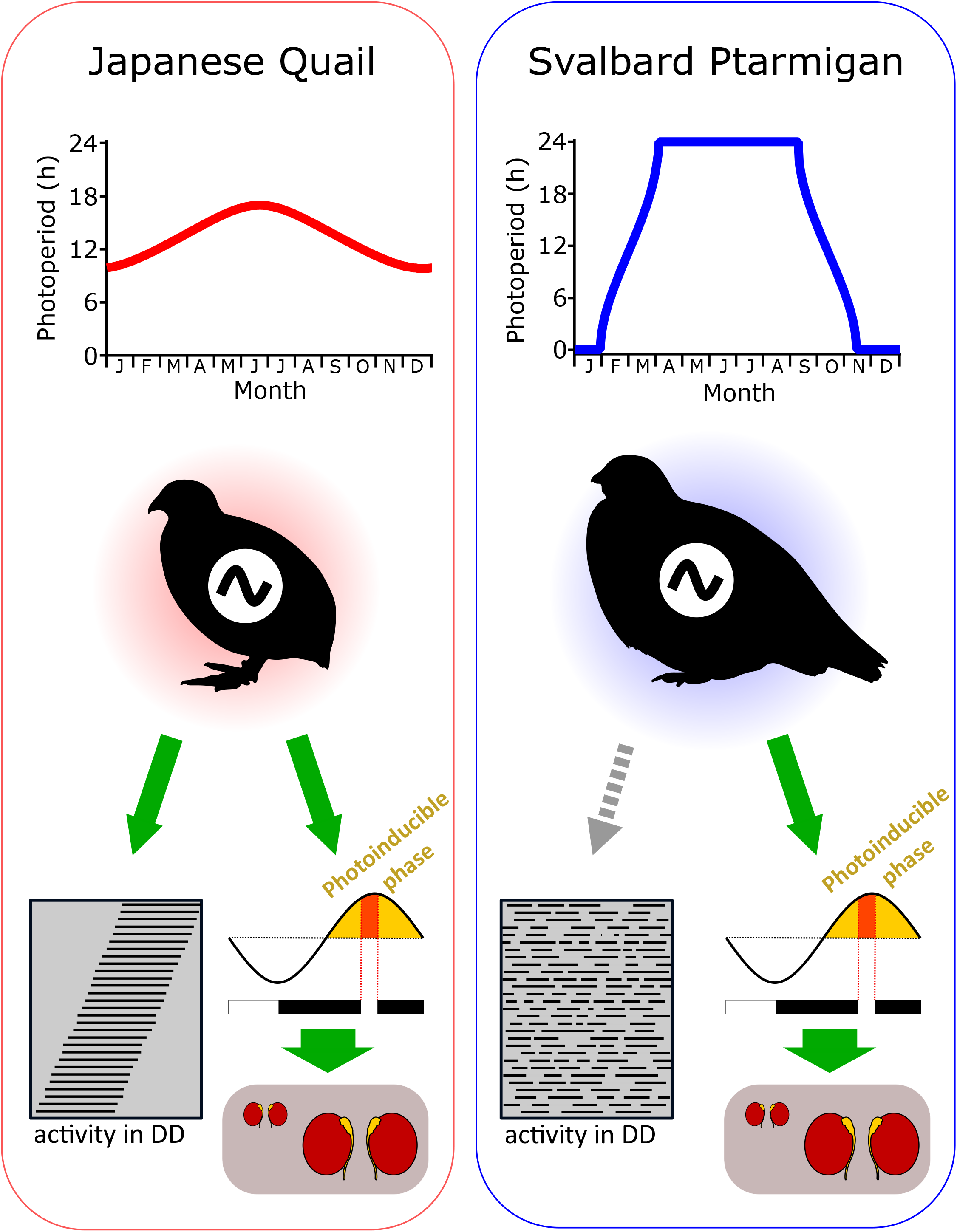
Adaptation of the circadian system to the Arctic. The Japanese Quail (left panel) uses its circadian system to control activity, as it retains a free running rhythm in prolonged constant darkness (DD) [40, 41]. The circadian system is also employed for photoperiodic time measurement. This is supported by studies using skeleton photoperiods that trigger long day responses, e.g. developing gonads, when the second light-period coincides with the photoinducible phase [10, 19, 42, 43]. We show here that its arctic relative, the Svalbard ptarmigan (right panel), retained its circadian system to sustain a rhythm of photosensitivity and responds to a correctly timed skeleton photoperiod in the same manner as the quail does. However, due to its high latitudinal environment and the special photic conditions of Svalbard we propose that the functional circadian system exhibits weak or no control over behavioural output.

## ACKNOWLEDGMENTS

We thank the animal technicians from the Arctic Chronobiology and Physiology research group: Hans Lian, Hans-Arne Solvang and Renate Thorvaldsen. Their experience and dedication is indispensable to our research. We would also like to thank Vebjørn J. Melum for his help with animal husbandry and Hugues Dardente for his help with the *in situ* hybridizations.

## AUTHOR CONTRIBUTION

Conceptualization, all; Methodology, all; Validation DA & GCW,; Formal analysis, DA & ACW; Investigation, all; Resources, GCW & DGH; Data curation, DA; Writing – Original draft, DA & ACW; Writing Review & Editing, DA, DGH & ACW; Visualization, DA, GCW & ACW; Supervision, DGH, GCW & ACW; Project Administration, all; Funding Acquisition, DGH;

## STAR⋆METHODS

### KEY RESOURCES TABLE

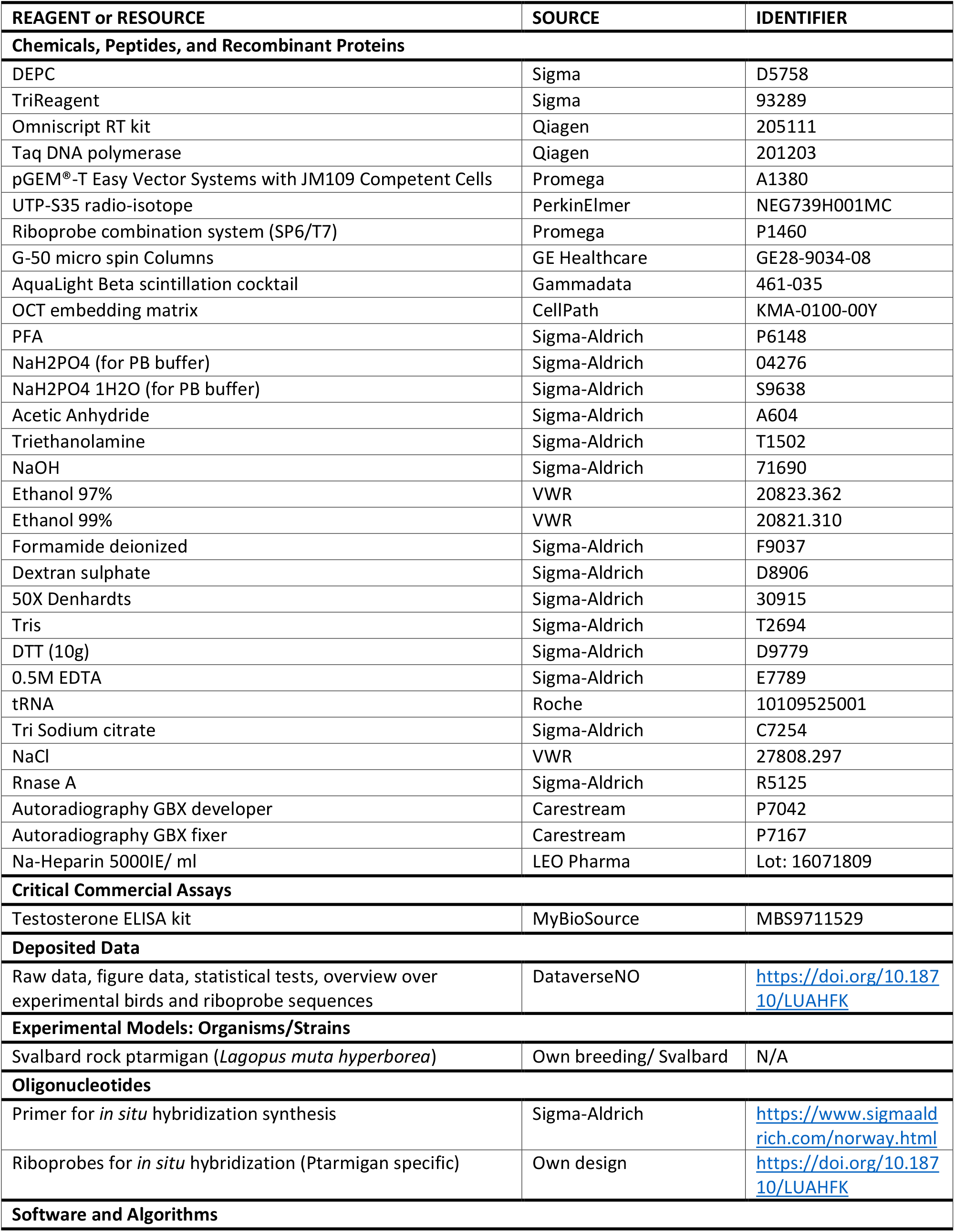
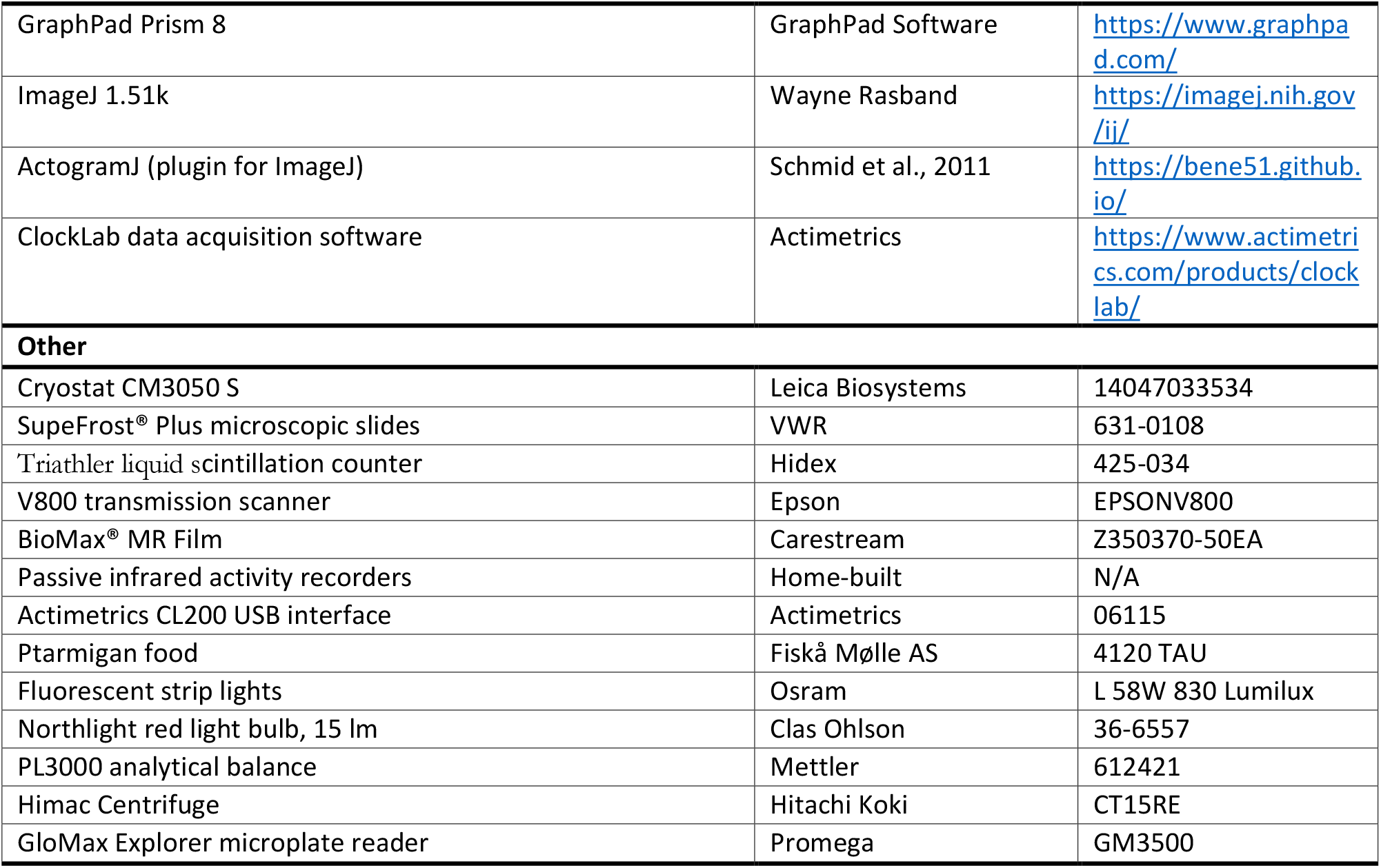

### RESOURCE AVAILABILITY

#### Lead contact

Further information and requests for resources and reagents should be directed to and will be fulfilled by the lead contacts, Alexander West or David Hazlerigg

#### Materials Availability

Ptarmigan specific *in situ* hybridization riboprobes generated for this study are available upon request.

#### Data and Code Availability

All material and data generated during this study are available at DataverseNO https://doi.org/10.18710/LUAHFK.

### EXPERIMENTAL MODEL AND SUBJECT DETAILS

This study was conducted on captive Svalbard ptarmigan (*Lagopus muta hyperborea*). The Svalbard ptarmigan is a subspecies of the Rock ptarmigan (*Lagopus muta*) and is a non-migratory and therefore permanent inhabitant of the high arctic archipelago of Svalbard (74 to 81 °N). Even though these birds are capable flyers they are isolated from other rock ptarmigan population, as reflected in their low genetic diversity [44] and the specificity of their phenotype compared to other ptarmigan populations, e.g. high amplitude body mass cycles [45].

Our facility located at the University of Tromsø operates a breeding program for Svalbard ptarmigan, which is regularly supplemented by birds caught in Svalbard. Experimental birds were hatched from eggs laid by captive Svalbard ptarmigan held in outside-aviaries. Hatching takes place between June and August in each breeding season and chicks are either raised outdoors on the ground or indoors at a photoperiod corresponding to the on- and offset of natural civil twilight in Tromsø (69° 39’N, 18° 57’E). Birds used for our study were transferred into individual cages in light and temperature controlled rooms in September 2017 for the circadian experiment (Figure 1) and in September 2018 for the skeleton photoperiod experiments (Figure 2 and 3). Birds of different sexes were housed together and each room held a maximum of twelve birds for the circadian experiment and a maximum of six bird for the skeleton photoperiod experiments. In both years, the initial photoperiod at transfer was L12:D12 which was thereafter gradually decreased to the respective photoperiodic treatments, which is L6:D18 for the circadian experiment and constant darkness (DD) for the skeleton photoperiod experiments. All birds were fed standardised protein food *ad libitum* (Fiskå Mølle) and provided with fresh water. Light was provided by fluorescent strip lights (Osram) delivering approximately 1000 lux at floor level. Permanent red light illumination (Clas Ohlson) was provided in order to allow husbandry in DD and outside the light hours.

Both sexes were used for the experiments as we have not seen any sex differences in hypothalamic gene expression in our previous study [3].Seasonal rhythm in body mass, activity and food intake is also similar between the sexes [5, 25]. A full table with all birds with their respective experimental group and their respective data is available online at DataverseNO (https://doi.org/10.18710/LUAHFK).

All animals were kept in accordance of the EU directive 201/63/EU under licences provided by the Norwegian Food Safety authority (Mattilsynet, FOTS 7971 for the circadian experiment, FOTS 14209 for the skeleton photoperiod experiments).

## METHOD DETAILS

### Circadian experiment (Figure 1)

Photoperiod was gradually decreased from September 2017 until reaching L6:D18 in mid-November 2017. The circadian experiment took place on the 21st and 22nd December 2017. The experimental birds were divided into two groups. The short photoperiod group (SP-group, n = 20) was kept under L6:D18 while the constant light group (LL-group, n = 16) was directly transferred from L6:D18 into LL on the 21st December. Both groups were then sampled five times with an interval of five hours. The sampling timed are given in Zeitgeber time (ZT) for the SP-group and CT for the LL-group and are as followed: ZT/ CT 3, 8, 13, 18 and 23 (ZT 0 corresponds to light-on switch for the SP-group). Birds were euthanized and brains were removed within five minutes. Brains were rapidly transferred onto a cooled metal block and ultimately stored at −80 °C until further processing. Brains from four birds were taken per sampling point. However, only three brains could be used for the CT 13 sampling point in the LL-group because one brain was damaged during the sampling procedure. ZT 3 was only sampled once and was used for plotting of both groups as there is no experimental difference between the groups at this point. ZT 3 was, however, excluded from the statistical analysis. All bird IDs and their corresponding sampling time points is available online at DataverseNO (https://doi.org/10.18710/LUAHFK).

### First skeleton-photoperiod experiment (Figure 2)

Photoperiod was gradually decreased from September 2018 until reaching DD (dim red light) on the 13th December 2018 in which they remained until the start of the experiment in the middle of January 2019. On the 19th January 2019 all birds were transferred into L6:D18. This marked the start of this experiment, which lasted 12 weeks. The birds were divided into three groups. The SP control group (n = 10) remained under SP throughout the whole experiment (SP-group). The increasing photoperiod group (n = 12) was subjected to a stepwise increase in photoperiod (IP-group). The light-period was extended by shifting the light-off switch by two hours each week until reaching LL in week 10. Thereafter birds of this group remained in LL for two more weeks until the end of the experiment in week 12. The skeleton photoperiod group (n = 12) was subjected to a night break protocol in which the initial photoperiodic treatment of L6:D18 was split into two blocks of light from week 2 onwards (SkP-group). The first light-period of four hours remained fixed whereas the second light-period of two hours was shifted weekly backward by two hours. In week 10, the moving block of light joined with the start of the fixed light-period. At this point the light-period was not shifted further and the photoperiod was effectively L6:D18 again, yet shifted forward by two hours compared to the SP-group. Thereafter birds of this group remained in L6:D18 for two more weeks until the end of the experiment in week 12.

Body mass and voluntary food intake of all birds was measured weekly with an analytical scale (Mettler). VFI was measured once a week from all birds by measuring food eaten within a 24 hours period. In addition blood was taken weekly from four males per group for plasma testosterone measurements (blood was not taken in week 11 and 12). Locomotor activity of all experimental birds was continuously recorded as movement per minute by passive infrared sensors (home-built), mounted on the cage doors. Data were collected by an Actimetrics CL200 USB interface coupled to a PC with the ClockLab data acquisition software version 2.61 (Actimetrics).

All bird IDs and their respective photoperiodic treatment is available online at DataverseNO (https://doi.org/10.18710/LUAHFK).

### Second skeleton photoperiod experiment (Figure 3)

The second skeleton photoperiod experiment was conducted with birds from the SP-group (n = 10) from the first skeleton photoperiod experiment. This experiment started on 4th April 2019. The birds were separated into two groups. The SP-group (n = 5) further remained under L6:D18 for eight weeks and where sampled at the end of the experiment. The SkP-group (n = 5) went through a similar shifting skeleton photoperiod as described in the previous experiment. However, the two hour light-period was only shifted until reaching ZT 14-16 in week 6 upon which point birds remained on L4:D10:L2:D8 for additional two weeks before they were sampled (ZT 0 corresponds to the lights-on switch from the fixed four hour light-period). All birds of this experiment were euthanised on week 8 between ZT 0.5 and ZT 1.5. After the euthanasia brains were removed and rapidly transferred onto a cooled metal block until ultimately stored at −80 °C.

Measurements of BM, VFI and activity and blood sampling was conducted in the same manner as in the first skeleton photoperiod experiment and all bird IDs with their respective photoperiodic treatment is available online at DataverseNO (https://doi.org/10.18710/LUAHFK).

### Hormone measurement

Blood was taken weekly from four male birds of every group. In the first skeleton experiment, four birds were sampled in each group and each week, except in DD and week 1, in which only a total of four birds were sampled. The data from DD and week 1 was used to plot all groups but was excluded from statistical analysis. In the second skeleton photoperiod experiment, two male birds were sampled in each group and each week. Up to 1 ml of blood was taken with heparinized (LEO Pharma) syringes and transferred into 1.5 ml Eppendorf tubes stored on ice. Within 30 minutes, the blood was centrifuged at 3.000 rpm at 4 °C for 15 minutes (Hitachi Koki). After centrifugation, the plasma was pipetted from the sample and transferred into 60 μl aliquots. The aliquots were frozen at −80 °C until further processing.

Plasma Testosterone concentration was measured with a competitive inhibition ELISA kit (MyBioSource) following the manufacture’s manual. Optical density was measured by a microplate reader (Promega) at 450 nm.

### cDNA cloning and *in situ* hybridization

Gene expression of seasonal and clock genes in the PT and MBH for the circadian and second skeleton photoperiod experiment was measured by radioactive *in situ* hybridization. All *in situ* hybridization probes (*Tshβ, Dio2, Dio3, Per2* and *Cry1*) are based on RNA extracted from Svalbard ptarmigan brain tissue and were designed using a Icelandic ptarmigan genome [46]. Brain cryo-sectioning, probe synthesis and *in situ* hybridization were performed as reported previously [3, 47] and are described in short as follows.

RNA from Svalbard ptarmigan brain samples was extracted using TriReagent (Sigma-Aldrich) and the extracted RNA was converted into cDNA using the Omniscript RT kit from Qiagen. Subsequent PCR was performed with primers (Sigma-Aldrich) based on the Icelandic rock ptarmigan genome and Taq DNA polymerase (Qiagen). Correct sized PCR products were extracted, cloned into pGEMT easy vectors (Promega), sequenced and verified against the reference genome. Riboprobe sequences are available online at DataverseNO (https://doi.org/10.18710/LUAHFK).

Vectors were linearized and transcribed with a Promega T7/ SP6 Riboprobe combination system in combination with a 35S-UTP isotope (PerkinElmer). Radioactively labelled riboprobes were subsequently purified with G-50 micro spin columns (GE healthcare) and incorporation of the radionucleotide into the riboprobe was measured as counts per minute by a liquid scintillation counter (Hidex, scintillation cocktail form Gammadata).

Frozen brains, which were cryo-sectioned (Leica Biosystems) and mounted unto pre-coated adhesion microscopic slides (VWR), were fixed in 4 % PFA (in 0.1 M PB) for 20 minutes on ice. Sections were rinsed twice with 0.1 M PB for 5 minutes after fixation. Next sections were acetylated with 3.75 % v/v of acetic anhydride in 0.1 M triethanolamine buffer (0.05 N NaOH). Slides were rinsed twice with 0.1 M PB for 5 minutes after acetylation, dehydrated with stepwise increasing ethanol solutions (50 %, 70 %, 96 %, 100 % for 3 minutes each) and dried under vacuum for at least 1 hour. Dried sections were hybridized overnight at 56°C with radioactively labelled riboprobe in hybridization buffer (50 % deionised formamide, 10 % dextran sulfate, 1 x Denhardt’s solution, 300 mM NaCl, 10 mM Tris, 10 mM DTT, 1 mM EDTA, 500 μg/ml tRNA). The amount of added riboprobe equals 10^6^ counts per minute for each microscopic slide. Hybridized sections were washed with 4 x saline sodium citrate (SSC) solutions (3 x 5 minutes) and treated with RNase-A solution (20 μg/ml RNase A, 500 mM NaCl, 1 mM Tris, 1 mM EDTA) for 30 minutes at 37 °C. After RNase-A treatment stringency washes were performed with SSC of decreasing concentration: 2 x SSC (2 x 5 minutes), 1 x SSC (1 x 10 minutes), 0.5 x SSC (1 x 10 minutes), 0.1 x SSC (30 minutes at 60°C), 0.1 x SSC (rinse). SSC solutions were each supplemented with 1 mM DTT.

After stringency washing slides were dehydrated in stepwise increasing ethanol solutions (50 %, 70 %, 96 %, 100 % for 3 minutes each) and dried under vacuum. Once sections were dry, they were exposed to autoradiographic films (Carestream) for 10 to 25 days. Exposed films were developed (Carestream), fixed (Carestream) and digitalised with an transmission scanner (Epson). Optical density (OD) was measured with ImageJ (Wayne Rasband).

## QUANTIFICATION AND STATISTICAL ANALYSIS

All graphs and statistical tests were prepared in GraphPad Prism (Version 8.3.0, San Diego, CA, USA). Seasonal and clock gene expression of the circadian experiment was analysed with 2-way ANOVA with *post hoc* Sidak’s multiple comparisons test (Figure 1D-E). 2-way ANOVA with *post hoc* Tukey’s multiple comparisons test was used to examine changes in body mass, activity (in activity/ day and in activity/ day divided by the photoperiod in h), plasma testosterone and food intake in the first skeleton-photoperiod experiment (Figure 2B-E and S1). Activity, body mass, food intake and plasma testosterone of the second skeleton photoperiod experiment was analysed by 2-way ANOVA with *post hoc* Sidak’s multiple comparison test (Figure S2). Relative gene expression between the SP-group and SkP-group of the second skeleton photoperiod experiment was tested with unpaired t-tests. Activity was normalized by dividing counts per minute of each bird by its 99^th^ percentile and actograms (Figure 2A, 3A and S3) were plotted with ActogramJ [48], a plugin for ImageJ (Wayne Rasband).

Results of statistical tests are available online at DataverseNO (https://doi.org/10.18710/LUAHFK).

## SUPPLEMENTAL INFORMATION

**Figure S1. Response in body mass and food intake to increasing and skeleton photoperiod.**

(A) Weekly body mass and is displayed as mean ± SEM

(B) Weekly voluntary food intake measured as grams of food eaten in a 24-h period. Data is presented as mean ± SEM.

**Figure S2. Physiological and endocrine responses in the second skeleton photoperiod experiment.**

(A) Activity measured as counts/ day and displayed as mean ± SEM

(B) Weekly body mass is displayed as mean ± SEM.

(C) Weekly body mass changes displayed as mean ± SEM.

(D) Weekly voluntary food intake measured as grams eaten in a 24-h period and displayed as mean ± SEM.

(E) Weekly plasma testosterone in male birds measured in ng/ml and displayed as mean ± SEM.

**Figure S3. Double plotted actograms of all experimental birds of the skeleton photoperiod experiments.**

(A) Actograms correspond to experimental design of Figure 2A

(B) Actograms correspond to experimental design of Figure 3A

