## Supplementary figures and images for "Adaptive value of circadian rhythms in High Arctic Svalbard ptarmigan"

### Supplemental Figure 1

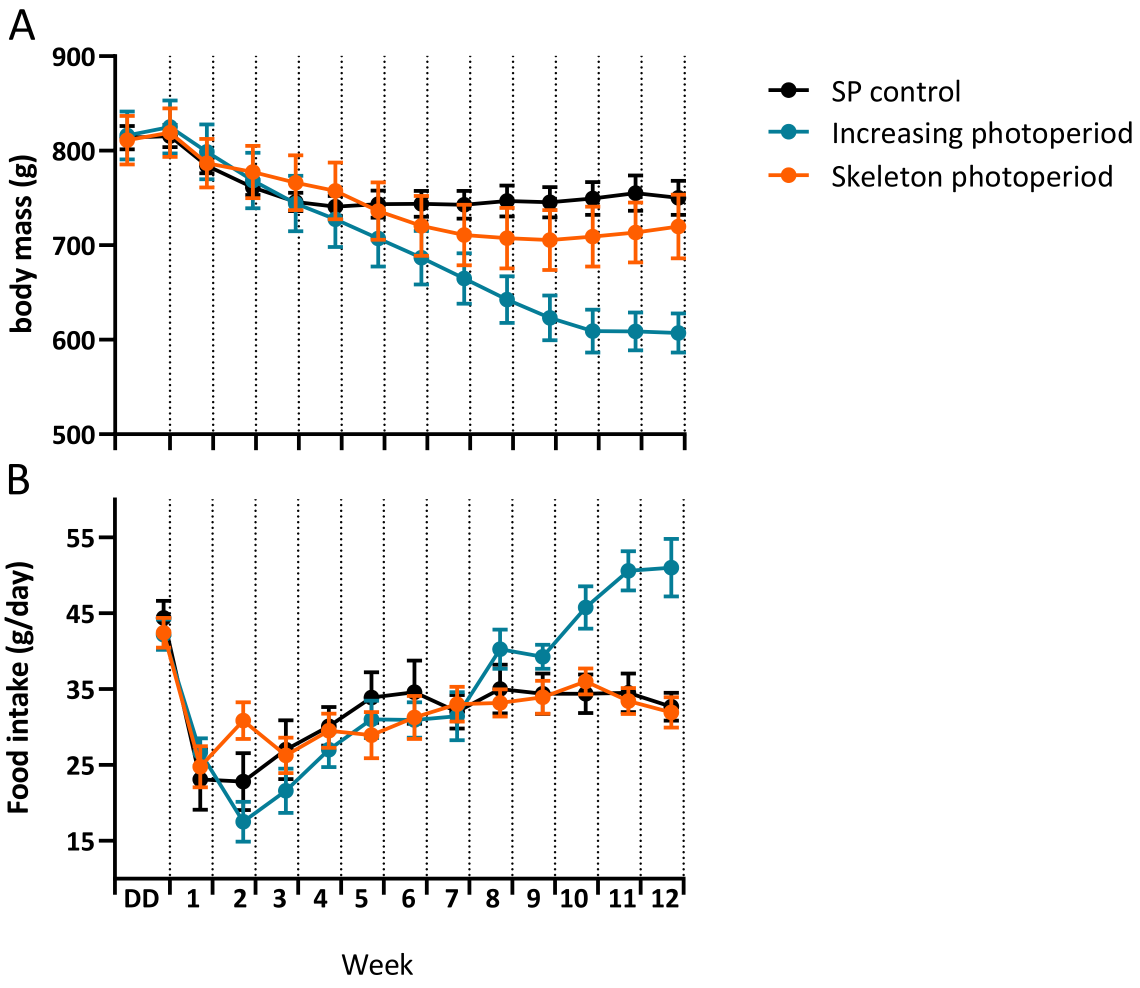

### Supplemental Figure 2

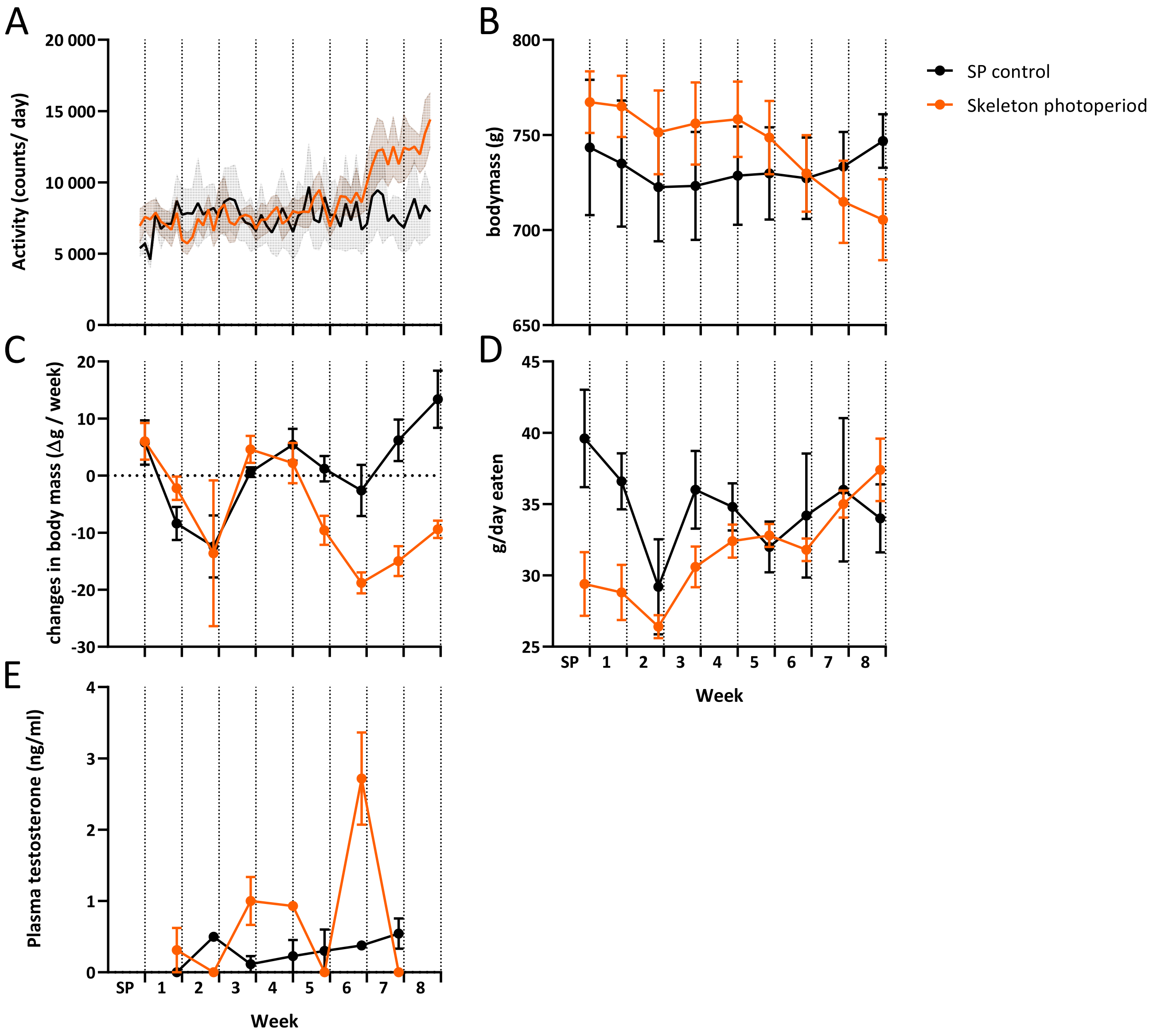

### Supplemental Figure 3

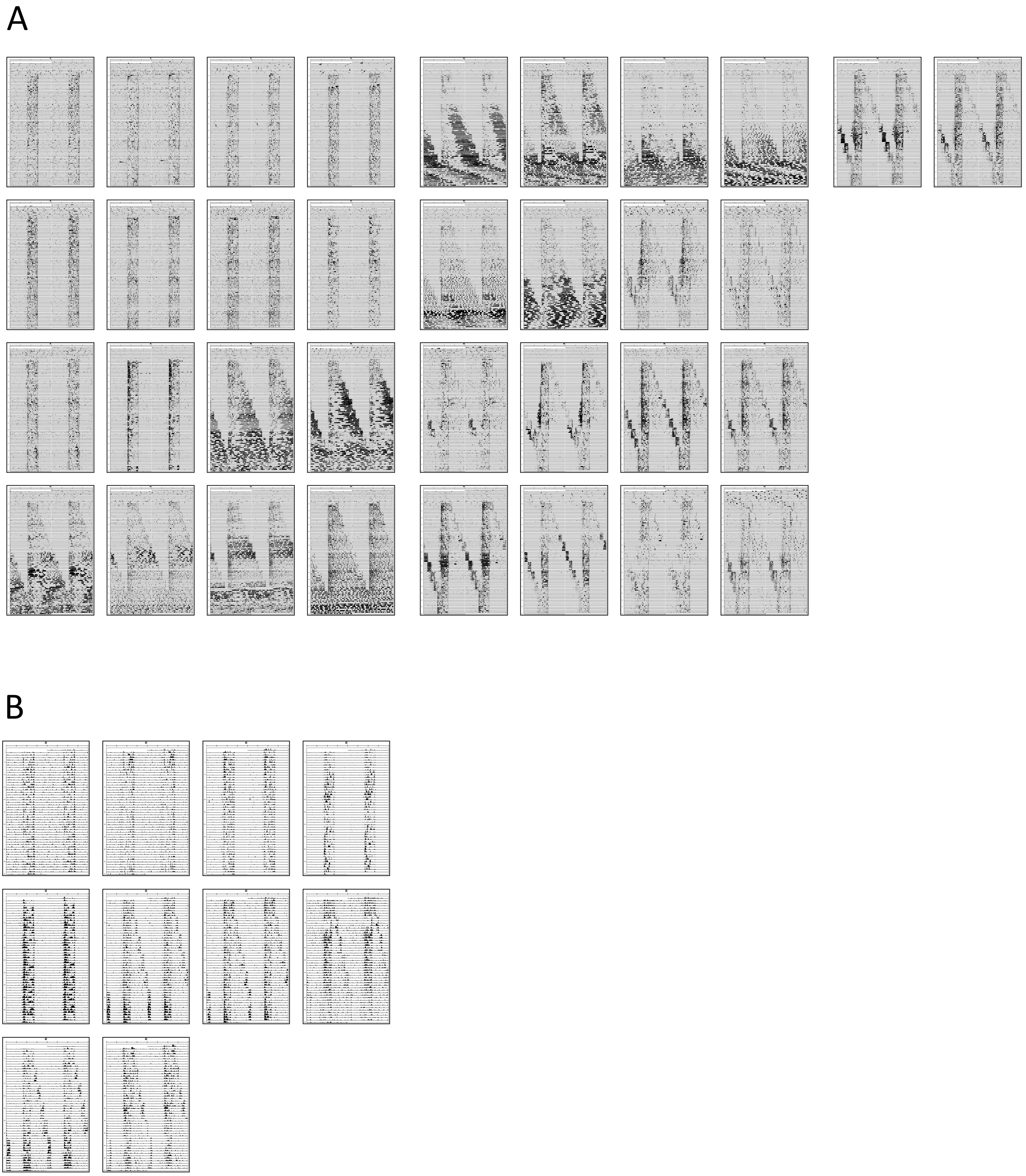


Figure S3.
